# The effect of a novel powerful ABA mimic on the improvement of color in grapes and its mechanism

**DOI:** 10.1101/2020.03.24.005371

**Authors:** Shanshan Ding, Chuanliang Che, Zhihong Xu, Xiaoying Du, Junkai Li, Jia-Qi Li, Yumei Xiao, Zhaohai Qin

## Abstract

Pigment content is an important quality attribute in the grape industry, and anthocyanins are the major fruit pigments. iso-PhABA is a novel and excellent ABA analog capable of the antimetabolic inactivation of ABA. In this study, we found that iso-PhABA improved the coloration of grape berries more obviously than ABA at concentrations of 2 mg/L and 5 mg/L in two grape varieties in China. iso-PhABA treatment enhanced the anthocyanin content in the two grape varieties; specifically, the anthocyanin and delphinidin contents increased in both the ‘Jufeng’ and ‘Xiahei’ varieties. An enzymatic activity test showed that iso-PhABA significantly promoted four key enzyme activities catalyzing anthocyanin biosynthesis. We also determined the affinity between iso-PhABA and ABA receptors using ABA as a control. The results indicated that iso-PhABA had significantly to moderately higher affinities for some ABA receptors, including PYR1, PYL2, PYL1, PYL3 and PYL10, which resulted in higher inhibition of the PP2C HAB1 in the presence of iso-PhABA than in the presence of ABA. iso-PhABA treatment increased the content of soluble sugars and grape yield without any apparent accompanying adverse effects on the quality of the grapes.

## Introduction

Grape is one of the main cultivated fruit varieties in China; according to the statistics of the Ministry of Agriculture and Rural Affairs of the People’s Republic of China, the area of viticulture in China has reached 11.98 million mu (799,000 hm^2^), and the output reached nearly 14 million tons at the end of 2015. The output of table grapes, including Jufeng, Xiahei, Fujisaka, Jingya and other European and American varieties, has ranked highest in the world since 2011 [1]. Jufeng (Kyoho), a tetraploid European-American hybrid derived from ‘V. labrusca × V. vinifera’ in Japan in 1937 [2], was introduced in China in 1959. Xiahei, a triploid seedless European-American hybrid grape, was derived from Vitis vinifera L. × Vitis labrusca L. in Japan in 1968 and introduced in China in 2000 [3]. These two table grape cultivars account for the majority of the market.

The commercial value of grapes is influenced by their appearance, including color. The color of grapes is mainly controlled by the content and type of anthocyanins [4-6]. High anthocyanin contents not only give grapes a beautiful appearance but also help to scavenge free radicals (ROS) effectively in the body [7] for the prevention of cardiovascular diseases [8,9] and other diseases. Thus, the production of grapes with beautiful colors has attracted increasing interestfrom both farmers and researchers. However, the coloration of grapes is greatly affected by environmental factors, such as high temperature [10]. In this situation, exogenous agents, especially plant growth regulators that help promote grape coloration [11], are the best choice for coloration improvement. In some areas, ethephon is a commonly used regulator that is used on red grapes to improve berry color; however, ethephon treatment can lead to berry softening, reducing the commercial value of the grapes [12]. Abscisic acid (ABA) is a natural plant endogenous hormone involved in the whole growth process of plants from seed germination to shedding [13]. It has long been known that ABA can promote grape coloration [14], but it is not commercially available because of its high cost. Recently, a new method for producing S-ABA by using gray mold organisms was developed [15,16], making it economically available for application, which greatly promotes the research and application of ABA on grapes and other fruits for color improvement [14,17-21]. However, due to the instability of ABA both in vivo and in vitro [22,23], the exogenous application of abscisic acid is somewhat limited. Over the past decades, numerous institutes, including our team, have focused on the development of new stable ABA analogs with better in vivo antimetabolite and in vitro anti-photo-isomerization activities [24-27]. In a previous study, we developed 2′,3′-iso-benzoabscisic acid (iso-PhABA), an excellent and easily prepared ABA analog that can inhibit metabolism in vivo [28]. This study mainly focused on the role of iso-PhABA in the pigmentation and quality of the two grape varieties ‘Jufeng’ and ‘Xiahei’ to discover a novel and potential colorant for grapes.

## Materials and methods

### Plant material and ABA/iso-PhABA treatments

This study was conducted in 2018 at experimental vineyards located in Hebei Province (36°01’N 113°04’E, China) and Hubei Province (29°01′N 108°21′E, China). Two major grape cultivars, ‘Jufeng’ and ‘Xiahei’, were used for the studies in Hebei and Hubei, respectively. The vineyards were managed under a conventional soil tillage management system. ABA solution was prepared with 1% S-ABA powder (Sichuan Lomon Fusheng Technology Co., Ltd.), and iso-PhABA solution was prepared with a homemade 1% soluble powder of iso-PhABA. A randomized block design was carried out with three blocks and five treatments, and each treatment in the block consisted of 3 individual vines. Based on our previous research (un-published data in 2017), three ABA and iso-PhABA concentrations (1, 2 and 5 mg/L) were used. During the early stages of grape veraison (20% - 25%), each of the chosen clusters was irrigated with 5 L of ABA or iso-PhABA solution with concentrations of 1, 2 or 5 mg/L. For all experiments, control clusters received no ABA treatment or iso-PhABA treatment, and there were three replicates for each treatment.

### Sampling

Grape berries in the vineyards were harvested randomly from the upper, middle and lower parts of different ears of the same grape at 0, 5, 10, and 15 days after treatment. The freshly harvested berries were selected on the base of similar size and the absence of physical injuries or insect infections. A total of 30 berries from each replicate were collected, washed with distilled water and precooled to 25 °C. A subsample of 10 berries was taken for measuring berry weight, and a similar subsample was used to determine the total soluble solid content, soluble sugar content, titratable acid content and tannin content. The remaining berries were dissected into seeds and skins, and skins were used to determine the anthocyanidin, cyanidin, and delphinidin contents and relative enzyme activity. The berries used for the measurement of the total soluble solid, soluble sugar, titratable acid, tannin, anthocyanidin, cyanidin, and delphinidin contents and the relative enzyme activity were immediately frozen in liquid nitrogen and stored at −80°C for subsequent analyses.

### Measurements of berry weight, total soluble solids, soluble sugar, titratable acid and tannin

Berry weight was measured with a balance (SJGD-JPT-1C, Beijing Heng Aode Instrument Co., Ltd.) in the field. Total soluble solids were measured with a pocket refractometer (PAL-2, Atago). Titratable acidity was measured with a pH-stat titration method using a titrator (AT-710M, Kem). The soluble sugar and tannin contents were determined with a microplate reader (Infinite F50, Tecan).

### Analysis of anthocyanidin, cyanidin, delphinidin and enzyme activity

To extract anthocyanidins, 10 freeze-dried berry skins were ground to a fine powder in liquid N_2_, and anthocyanins were extracted according to the methods of Cheng et al. [29] and Zoratti et al. [30]. Grape skin powder (1 g) was extracted with 20 mL of a 1 M HCl/methanol/water (1:80:19, v/v/v) mixture. The extraction was performed under ultrasound for 10 min at 100 Hz, followed by shaking in the dark at 25 °C for 30 min at a rate of 150 rpm. Then, the sample was centrifuged using a high-speed refrigerated centrifuge for 10 min at 10,000 rpm at 4 °C, and the supernatant was collected. The residues were re-extracted four times. All supernatants were collected and combined. The total anthocyanin content was measured at 530 nm using a microplate reader (Infinite F50, Tecan), and the contents of cyanidin and delphinidin, the components of anthocyanidins, were also measured using a microplate reader (Infinite F50, Tecan). The activities of PAL, CHI, DFR and flavonoid glycosyltrans-ferase (UFGT) were evaluated as per the protocols of ELISA Kit (Beijing gersion Bio-Technology Co., Ltd) and calculated with a microplate reader (Infinite F50, Tecan).

### Determination of affinity (K_d_) with ABA receptor and inhibitory activity against phosphatase (HAB1)

The affinity between small molecules and receptors was determined by the microthermal swimming method to evaluate the affinity (K_d_) between small molecules and proteins. The receptor protein was diluted to 10 μM, and the receptor protein was marked according to the standard protocol of the Nano Temper manual. The small molecules of different concentrations were mixed with the receptor in Tris buffer and incubated at room temperature for 5 min. The samples were drawn with the Nano Temp standard capillary and placed in an MST instrument for testing. The experiments in each group were repeated three times.

The activity of phosphatase was determined by the Serine/Threonine Phosphatase Assay System kit from Promega Company. The PYR/PYL/ RCARs with a final concentration of 3 μM and 1 μM HAB1 were dissolved in Tris buffer. After incubation at room temperature for 20 min, a small molecule was measured with a final concentration of 10 μM and a polypeptide substrate of 5 μM. After incubation at room temperature for 20 min, 50 μL molybdate pigments/additives were added to terminate the reaction. After half an hour, the absorbance value of the system at 630 nm was determined by an enzyme labeling instrument. The phosphatase activity of HAB1 was calculated according to the formula. The experiments in each group were repeated three times.

### Statistical analysis

All experiments were carried out in triplicate, and the results were expressed as the mean ± standard deviation (SD). Statistical procedures were performed with SPSS version 19.0 statistical package (SPSS Inc., Chicago, IL, USA). Data from the different experiments were subjected to analysis of variance (ANOVA). Significant differences between means were determined by use of Tukey’s test at p < 0.05.

## Results and discussion

### iso-PhABA improves the coloration of grape berries

To confirm the role of iso-PhABA in grape skin coloration, experiments were conducted with grape varieties ‘Jufeng’ and ‘Xiahei’ in Hebei Province (North China) and Hubei Province (South China) in 2018. As shown in Figure 1, after 25% of the grapes were colored, the roots were treated with water, ABA and iso-PhABA. Similar results were observed in both Jufeng and Xiahei: the fruit coloration time after ABA and iso-PhABA treatment occurred earlier, and the color of the berries after ABA and iso-PhABA treatment was significantly deeper compared to that after water treatment for 10 days, and the color of the berries after iso-PhABA treatment was more obvious than that after ABA treatment at 2 mg/L and 5 mg/L, respectively. After treatment for 15 days, the coloration depth of the two types of grape peels in the 2 mg/L iso-PhABA treatment group was the darkest and was nearly the same as that of the 5 mg/L iso-PhABA treatment group, followed by the ABA treatment group and the water treatment group with the least coloration.

**Figure 1.**
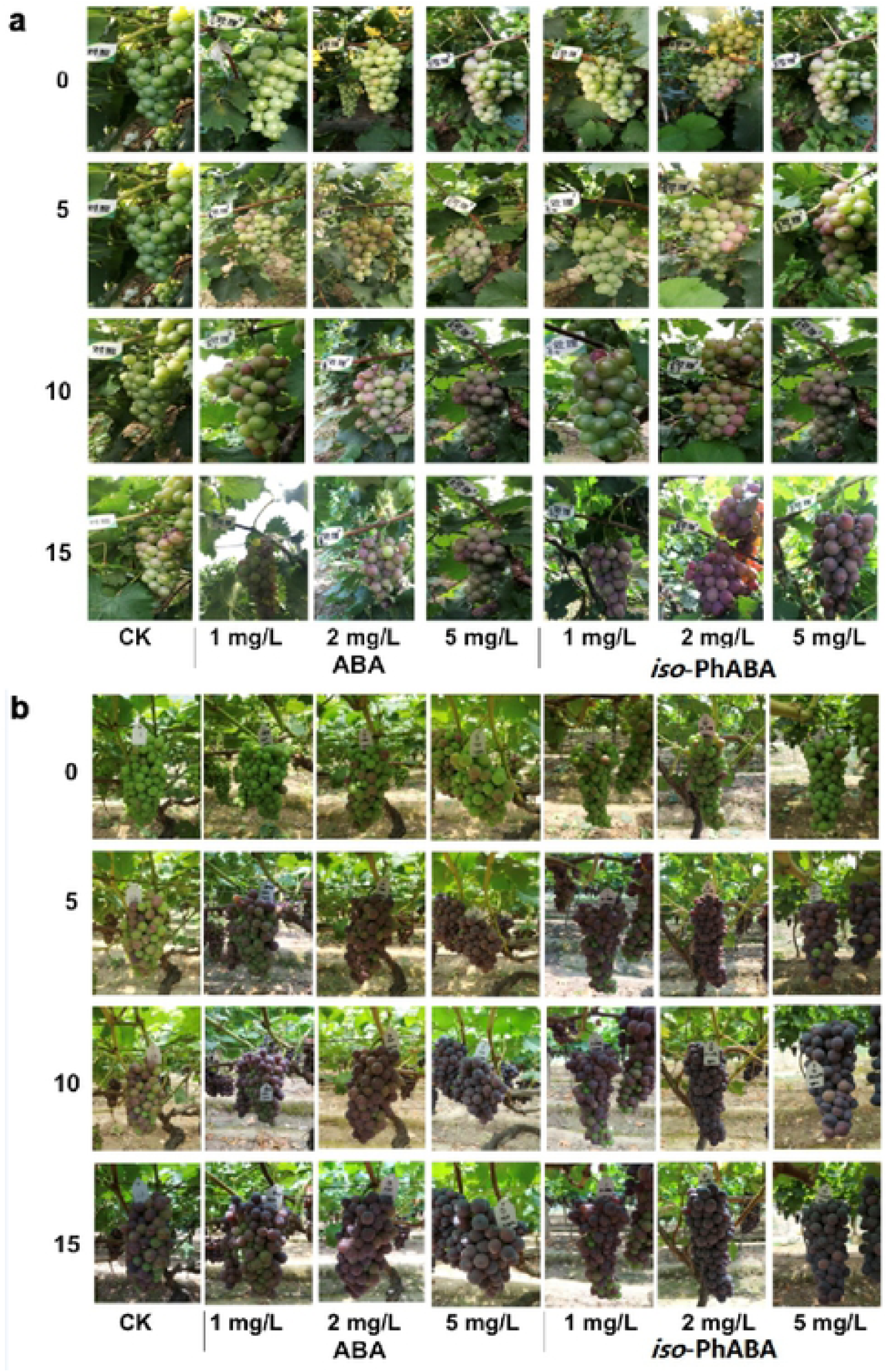
Coloration of grape skin after ABA and iso-PhABA treatment. (a. Hebei, Jufeng; b. Hubei, Xiahei)

### iso-PhABA improves the anthocyanin content in grape berries

The anthocyanin content determines the color of grape skin, and a higher content provides darker coloration. According to the above results, we determined the total anthocyanin content in grape skin. As shown in Figure 2, after treatment for 5 days, the anthocyanin content began to increase significantly. After 10 days, the cyanine content in ‘Jufeng’ and ‘Xiahei’ grape berries treated with 1 mg/L ABA was 5.38 ± 0.68 mg/g and 26.89 ± 3.42 mg/g, respectively; the contents were 6.22 ± 0.47 mg/g and 31.12 ± 2.36 mg/g under treatment with 1 mg/L iso-PhABA in ‘Jufeng’ and ‘Xiahei’ grape berries, respectively, while the contents in the control were 4.94 ± 1.48 mg/g and 19.01 ± 2.57 mg/g, respectively. On day 15, 1 mg/L iso-PhABA treatment resulted in a slightly higher cyanine content in ‘Jufeng’ and ‘Xiahei’ grape berries. With increasing concentrations of ABA and iso-PhABA, the cyanine content in grape berries increased. In the ‘Jufeng’ variety, the anthocyanin content at 2 mg/L and 5 mg/L iso-PhABA was obviously higher than that of the ABA treatment. This finding was consistent with the coloration depth trend, which was also consistent with previous studies of the effect of ABA on the content of cyanine in grapes [31]. Anthocyanins are composed of different forms, including cyanins and delphinidins.

**Figure 2.**
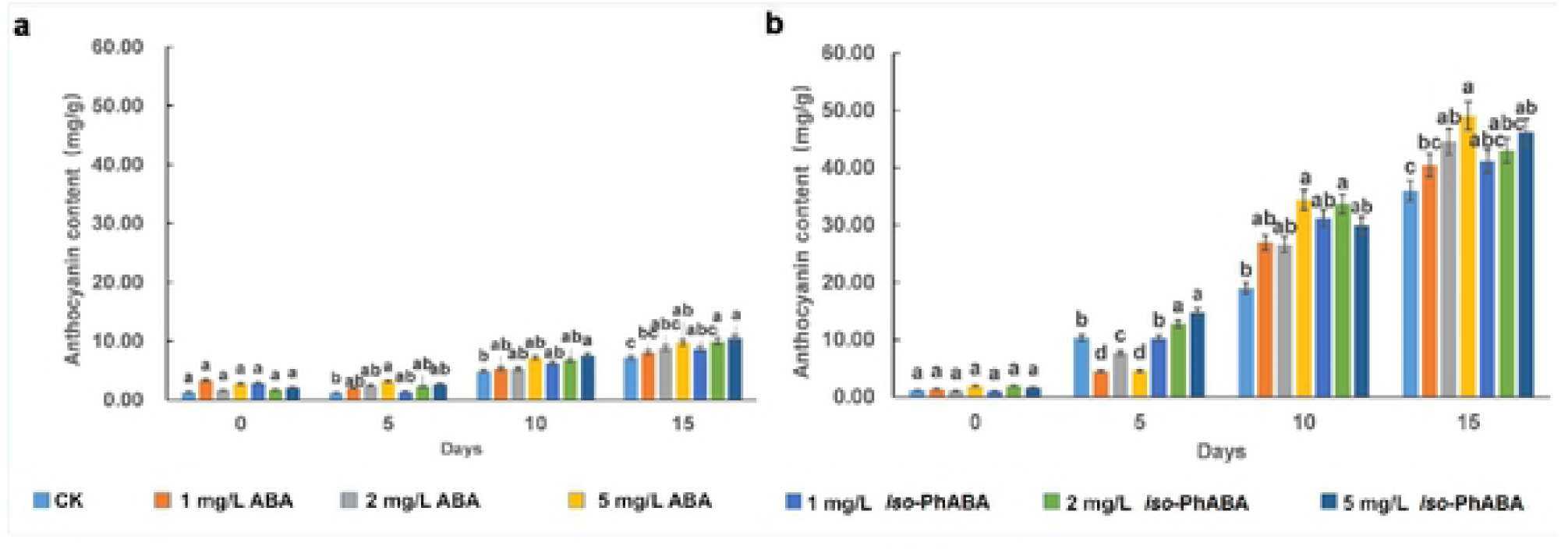
Anthocyanin contents in grape fruits treated with ABA and iso-PhABA. (a) Anthocyanin content in grape variety ‘Jufeng’ after treatment with ABA and iso-PhABA (planted in Hebei Province); (b) Anthocyanin content in grape variety ‘Xiahei’ after treatment with ABA and iso-PhABA (planted in Hubei Province).

We also analyzed the effects of ABA and iso-PhABA on the cyanin and delphinidin contents. The cyanin and delphinidin contents increased after ABA or iso-PhABA treatment (Figure 3). In ‘Jufeng’ grapes, the contents of delphinidin (ranging from 3.86 to 5.78 mg/g) and anthocyanin (ranging from 6.67 to 8.56 mg/g) in the ABA or iso-PhABA treatment groups were higher than those in the control group (3.30 mg/g and 5.80 mg/g) on the 5th day. Until day 15, the contents of anthocyanin and delphinidin in the ABA or iso-PhABA treatment were consistently higher than those in the control, and higher treatment concentrations resulted in higher anthocyanin and delphinidin contents. The highest contents of anthocyanin and delphinidin at day 15 were 7.14 ± 0.09 mg/g and 10.46 ± 0.10 mg/g (treatment with 5 mg/L iso-PhABA), respectively. However, in ‘Xiahei’ grapes, after ABA treatment, the content of delphinidin did not change significantly, and only the content of anthocyanin increased significantly. This indicated that ABA and iso-PhABA may affect the content of different kinds of anthocyanin monomers in different grape cultivars, which was consistent with previous findings [32,33].

**Figure 3.**
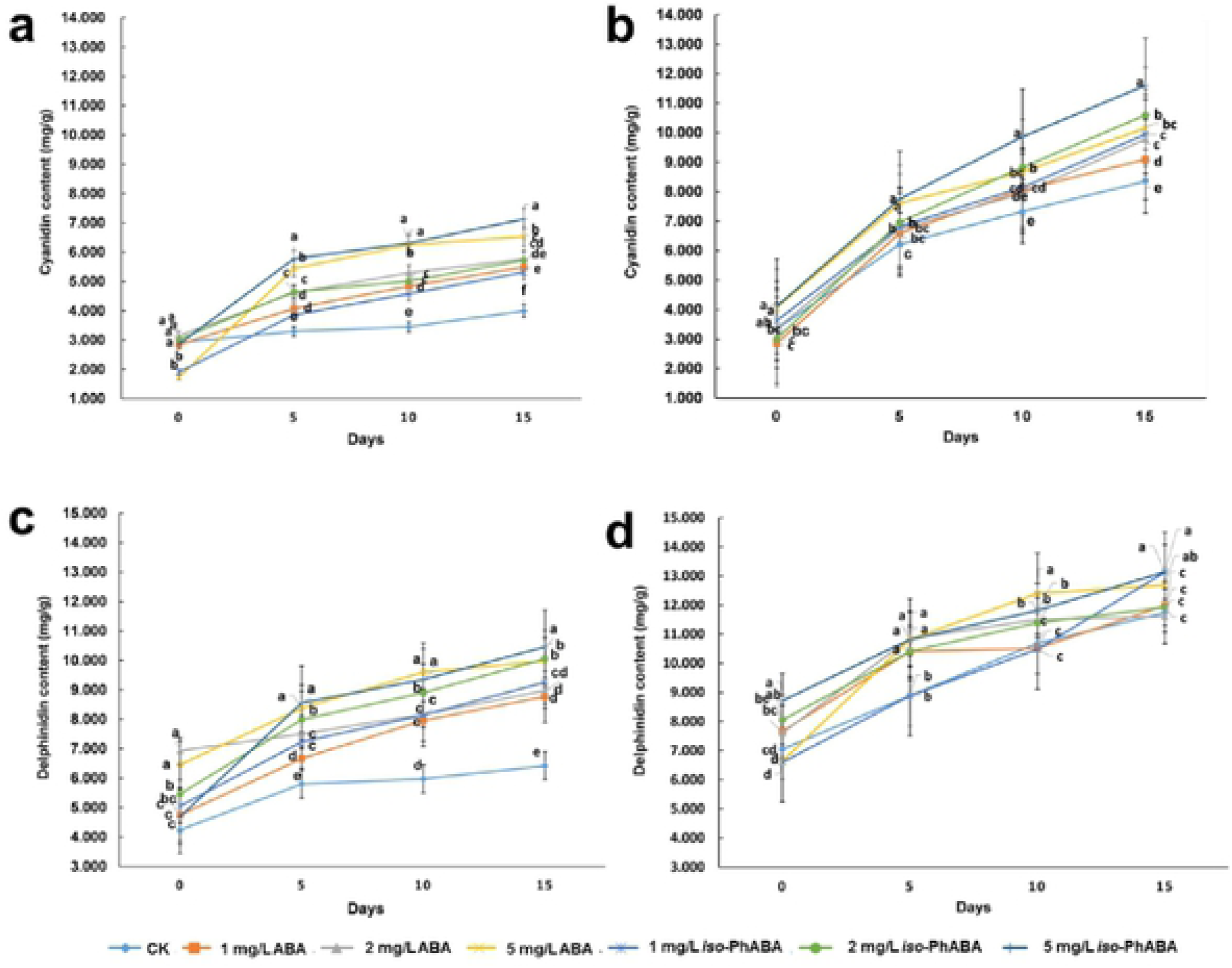
The content of cyanins and delphinidin in grape fruit after treatment with ABA or iso-PhABA. (a) The content of cyanidin in grape variety ‘Jufeng’ and (b) in ‘Xiahei’. (c) The content of delphinidin in grape variety ‘Jufeng’ and (d) in ‘Xiahei’.

### iso-PhABA enhances the activity of key enzymes involved in anthocyanin synthesis

Anthocyanins are synthesized in the cytoplasm of epidermal cells and accumulate in the skin of grape berries during their initial formation stages in vacuoles via a series of transport pathways [34]. The biosynthesis of anthocyanins is carried out via a series of catalytic reactions by enzymes, involving more than 20 steps throughout the whole process. The synthesis pathway of anthocyanins is mainly divided into three stages. The first stage is the phenylalanine metabolic pathway, which converts phenylalanine to coumaroyl CoA using the enzymes phenylalanine lyase (PAL), cinnamic acid hydroxylase (C4H) and coumaroyl CoA ligase (4CL). These enzymes are shared by many secondary metabolisms. The second stage is the flavonoid pathway, which converts coumarin CoA to flavonoids. This pathway is mainly carried out by chalcone synthase (CHS), chalcone isomerase (CHI), alkanone-3-hydroxylase (F3H), and dihydroflavonol 4-reductase (DFR) and is completed by the catalysis of leuco-feline dioxygenases (LDOXs). After modification by methylation and acetylation, anthocyanins of different structures are formed during the final stage [35]. Anthocyanins in grapes are mainly composed of methylated cyanine (red) and methylated delphinidin (blue-purple). The ratio of various anthocyanins and the level of their accumulation differ among different red, purple and black grape berries [36,37].

To further understand the mechanism of the effect of iso-PhABA on anthocyanin synthesis, we analyzed the activities of 4 key enzymes, PAL, CHI, DFR, and UFGT, in the synthesis of anthocyanins. The results are shown in Figure 4. The activities of the four enzymes in the ABA and iso-PhABA treatment groups were significantly higher than those in the control, and the activities of PAL, CHI and UGFT increased gradually with increased treatment time. In ‘Jufeng’, the activities of the enzymes decreased according to the ABA or iso-PhABA concentration as follows: 5 mg/L > 2 mg/L > 1 mg/L. The activities of PAL in ‘Jufeng’ grapes treated with 5 mg/L ABA and 5 mg/L iso-PhABA were 0.187 ± 0.002 U/g and 0.186 ± 0.002 U/g, which were higher than those of the control (0.130 ± 0.001 U/g) and other treatments. The activities of CHI in ‘Jufeng’ grapes treated with 5 mg/L ABA and 5 mg/L iso-PhABA were 3.685 ± 0.031 U/g and 3.617 ± 0.040 U/g, which were higher than those of the control (2.745 ± 0.034 U/g) and other treatments. The enzyme activities of DFR and UFGT in grapes treated with 5 mg/L ABA were 7319.33 ± 88.61 U/g and 6664.33 ± 111.34 U/g, and the activities of DFR and UFGT in grapes treated with 5 mg/L iso-PhABA were 7500.00 ± 37.64 U/g and 6507.33 ± 81.98 U/g. In ‘Xiahei’, the 5 mg/L iso-PhABA treatment significantly increased the activity of the 4 enzymes. The PAL activity in the 5 mg/L iso-PhABA treatment was 0.139 ± 0.005 U/g, equal to the PAL activity of the 5 mg/L ABA treatment (0.139 ± 0.005 U/g) and higher than that of the control (0.126 ± 0.002 U/g). The CHI activity of the 5 mg/L iso-PhABA treatment was 4.597 ± 0.026 U/g, which was significantly higher than that of the control (3.731 ± 0.049 U/g). The DFR activity of the 5 mg/L iso-PhABA treatment was 7313.33 ± 167.53 U/g, which was significantly higher than that of the control (5933.67 ± 51.48 U/g). The UFGT activity of the 5 mg/L iso-PhABA treatment was 10510.33 ± 160.35 U/g, which was significantly higher than that of the control (8745.67 ± 132.67 U/g). These results indicated that the effect of iso-PhABA was similar to that of ABA, which increases the activity of anthocyanin synthase and promotes the synthesis of anthocyanins. However, differences in the enzyme activities were observed between the two grape varieties, especially the activity of the PAL enzyme. Low concentrations (1 mg/L and 2 mg/L) of ABA or iso-PhABA did not significantly increase the activity of PAL in ‘Xiahei’ grapes, in contrast to the obvious changes in ‘Jufeng’ grapes when treated with low concentrations (1 mg/L and 2 mg/L) of ABA or iso-PhABA. We also found that the 5 mg/L ABA treatment had more obvious effects on PAL, DFR and UFGT activities in ‘Xiahei’ grapes, while in ‘Jufeng’ grapes, the 5 μg/mL ABA treatment had obvious effects on PAL, DFR, UFGT and CHI activities. These changes may be related to different environmental factors, such as temperature and light, in different provinces and the genetics of grape.

**Figure 4.**
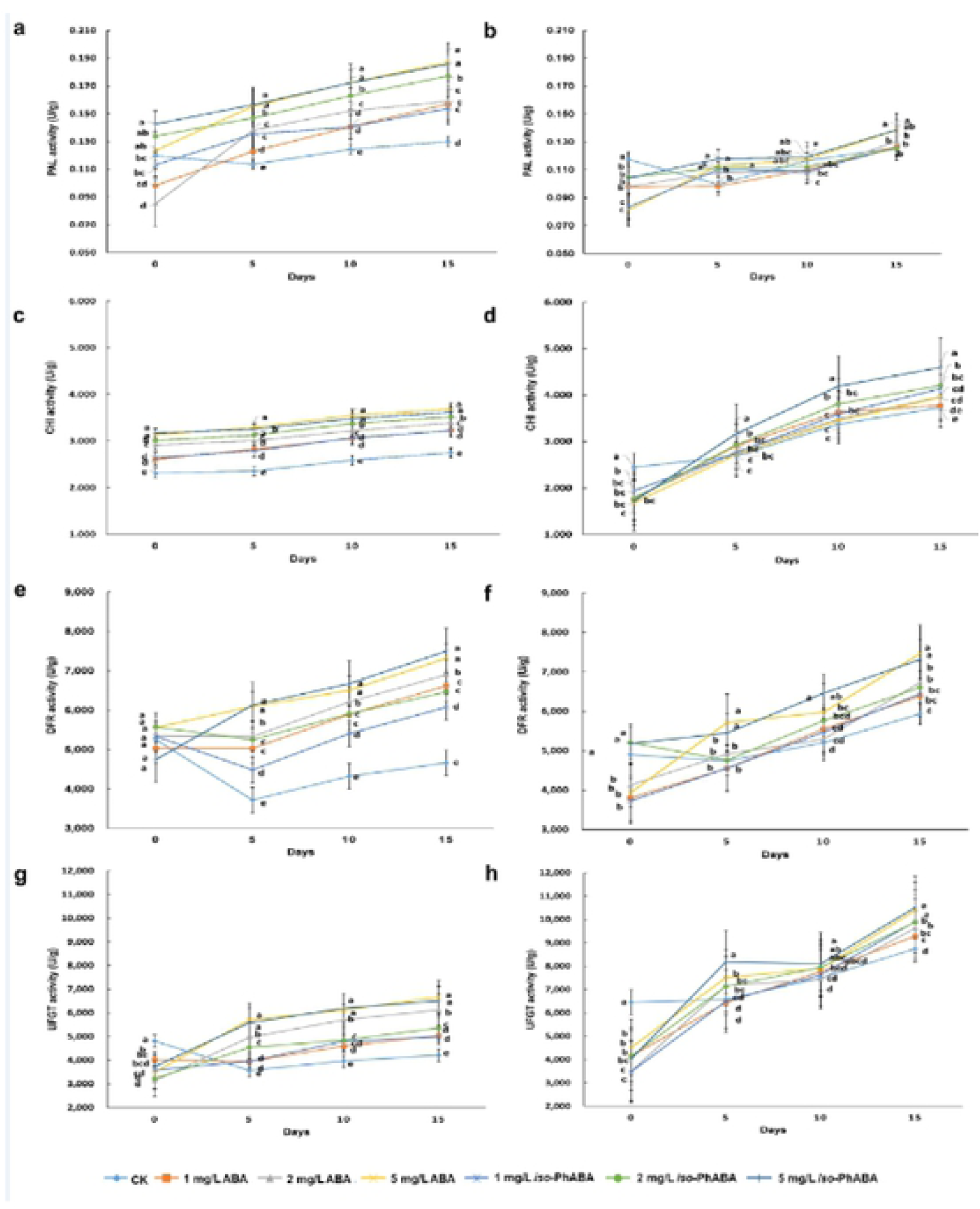
Activity changes of anthocyanin synthase after treated with ABA or iso-PhABA. The PAL activity analysis in grape variety ‘Jufeng’ (a) and ‘Xiahei’ (b) after treated with ABA and iso-PhABA; The CHI activity analysis in grape variety ‘Jufeng’ (c) and ‘Xiahei’ (d) after treated with ABA and iso-PhABA; The DFR activity analysis in grape variety ‘Jufeng’ (e) and ‘Xiahei’ (f) after treated with ABA and iso-PhABA; The UFGT activity analysis in grape variety ‘Jufeng’ (g) and ‘Xiahei’ (h) after treated with ABA and iso-PhABA.

### iso-PhABA can effectively activate the ABA signaling pathway

To better explain the mechanism by which the effect of iso-PhABA on the partial enzyme in promoting grape transformation is superior to that of ABA, we further explored the binding affinity of iso-PhABA with the ABA receptor and the inhibitory activity on the phosphatase HAB1.

The binding dissociation constant Kd value of the representative compound and some ABA receptors was determined by the MST experiment (Table 1). The Kd value is the equilibrium constant of the dissociation of the binary complex during the binding of the drug molecule to the receptor (binding site). Inversely proportional to the affinity between the two, the smaller the Kd value, the greater the affinity between the drug molecule and the receptor. The Kd value is affected by hydrogen bonds, electrostatic interactions, hydrophobicity, van der Waals forces and steric resistance. The results show that iso-PhABA and ABA showed strong binding forces with dimer and monomer receptors. The affinities of iso-PhABA and PYR1, PYL1, PYL2, PYL3 and PYL10 are better than those of ABA.

**Table 1.**
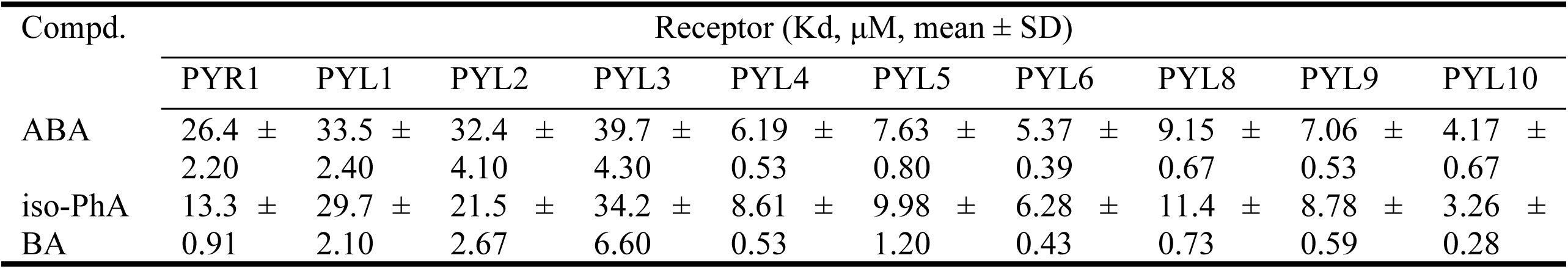
Comparison of ligand-receptor affinity between ABA and iso-PhABA.

After ABA binds to receptors, the conformations of the gate loop and latch loop change from open to closed, and then they can bind to PP2Cs and inhibit their activity. To evaluate the activation of the iso-PhABA signaling pathway at a deeper level, we measured the inhibitory effect of iso-PhABA on phosphatase HAB1 after binding to the ABA receptor. Table 2 shows that the binding of ABA and iso-PhABA with PYR1, PYL4 and PYL5 can almost completely inhibit the activity of HAB1. The inhibition activity of iso-PhABA on the phosphatase is slightly stronger than that of ABA, which indicates that iso-ABA is a complete ABA receptor agonist that triggers a series of signals downstream.

**Table 2.** Comparison of inhibitory activities of ABA and iso-PhABA after combination with PYLs on phosphatase HAB1. (% inhibition rate, ligand: PYLs: HAB1 = 10 μM: 3 μM: 1 μM)

| Compd. | PYR1 | PYL1 | PYL2 | PYL3 | PYL4 | PYL5 | PYL6 | PYL8 | PYL9 | PYL10 |
| --- | --- | --- | --- | --- | --- | --- | --- | --- | --- | --- |
| ABA | 99.56 | 99.64 | 98.13 | 98.29 | 99.24 | 98.23 | 99.62 | 98.72 | 99.03 | 99.63 |
| iso-PhABA | 99.75 | 99.19 | 99.63 | 98.95 | 99.51 | 98.95 | 98.86 | 97.69 | 99.15 | 99.83 |

### iso-PhABA treatment increased the soluble sugar content in grape berries

The effect of iso-PhABA on the quality of grapes was also evaluated in this study. First, the soluble sugar content in the grape berries was analyzed. As shown in Figure 5, compared to the control, ABA and iso-PhABA treatments increased the soluble sugar content. The soluble sugar content is related to the concentration of ABA or iso-PhABA, which was highest in the 5 mg/L ABA and iso-PhABA treatment groups, followed by the 2 mg/L treatment group, 1 mg/L treatment group, and finally the control. For example, in ‘Jufeng’ grapes, the content of soluble sugar treated with 5 mg/L iso-PhABA was 17.65 ± 0.79 μg/g, which was significantly higher than that of the control (11.33 ± 1.72 μg/g) on the 15th day. This result was consistent with other studies [12,33,38], indicating that ABA improved the soluble sugar content. It has been widely shown that the coloration of the fruit depends on the anthocyanin content, and high anthocyanin levels lead to good coloration. Anthocyanins are composed of anthocyanidins and sugar groups. Therefore, grape coloring is closely related to the sugar content, and only when the grape soluble sugar content reaches a certain concentration do grapes begin to color. This may explain why ABA and iso-PhABA improved the soluble sugar content.

**Figure 5.**
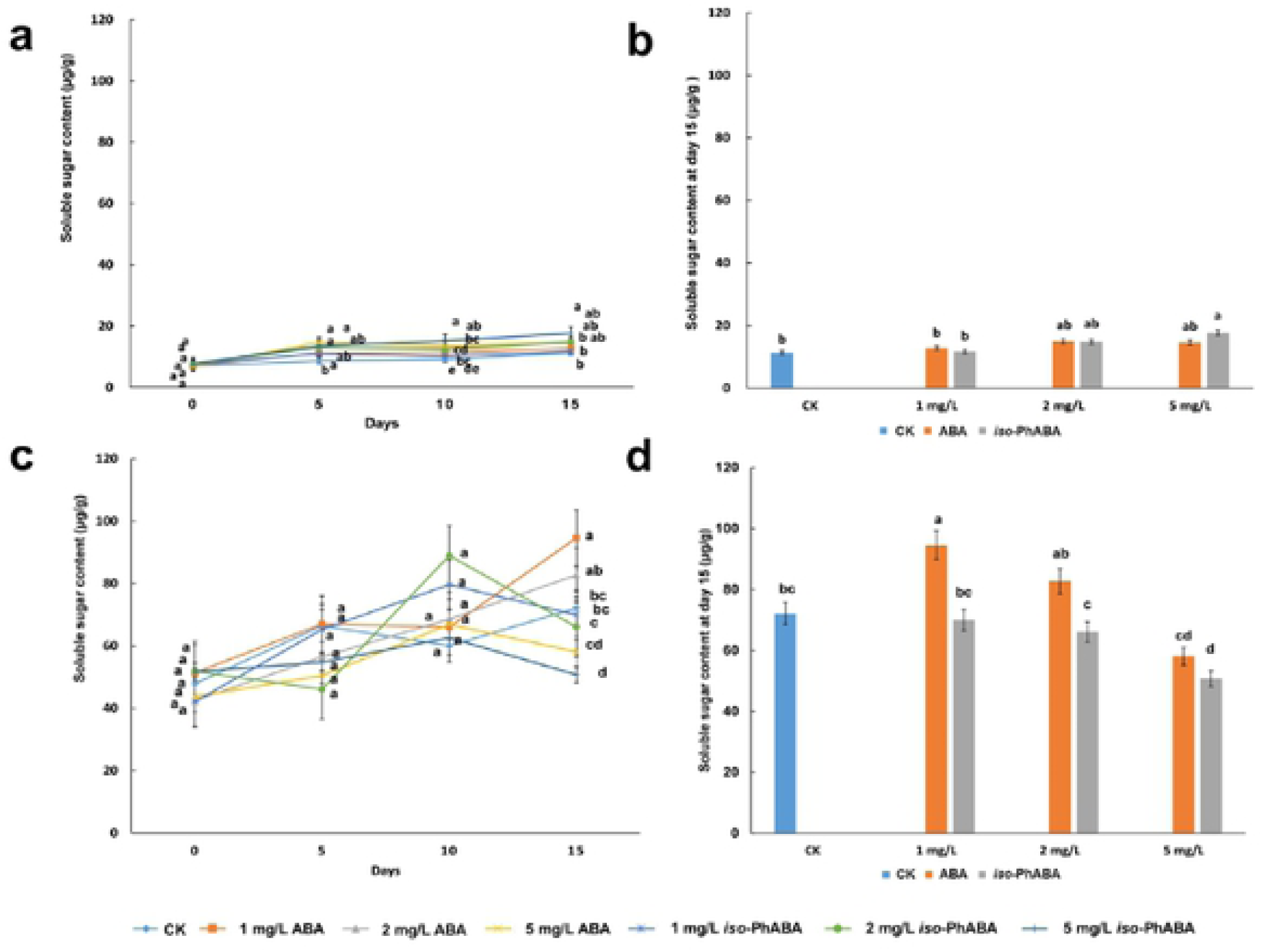
Soluble sugar content in grape varietes ‘Jufeng’ and ‘Xiahei’ treated with ABA and iso-PhABA. Soluble sugar content in grape varietes ‘Jufeng’ (a) and ‘Xiahei’ (c) treated with ABA and iso-PhABA; Soluble sugar content in grape varietes ‘Jufeng’ (b) and ‘Xiahei’ (d) after treated with ABA and iso-PhABA for 15 days.

Interestingly, we found that the accumulation of soluble sugars in the untreated group of ‘Jufeng’ grapes reached 11.33 μg/g (11.33 ± 1.72 μg/g) at day 15, while in the ABA- or iso-PhABA-treated group, the soluble sugar content reached 11.33 μg/g at day 10. A similar trend was detected in ‘Xiahei’ grapes. The accumulation of soluble sugar in the ABA- or iso-PhABA-treated group was faster than that of the control group, reaching 72.12 μg/g at day 10 or before day 15. This indicated that iso-PhABA promoted the ripening of the grape fruits, and we could harvest the grape berries 5 days in advance after treatment with iso-PhABA to fulfill market demands.

### Effects of iso-PhABA on grape yield and other qualities

To promote its application in agricultural production, it is necessary to evaluate the effect of iso-PhABA on grape yield and quality. Therefore, we further analyzed the weight of a single berry, soluble solid content, tannin content and soluble acid content in grape berries. As shown in Figure 6A, the single berry weight of the control increased with time, from 9.44 g to 10.84 g in ‘Jufeng’ grapes, and the single berry weights of the berries in other treatments were not significantly different from those of the control. In ‘Xiahei’ grapes, the single berry weight of the control increased from 3.21 g to 6.26 g, and ABA treatment and iso-PhABA treatment did not affect the single berry weight of grapes (Figure 6b). The soluble acid content was only determined for the ‘Jufeng’ variety, and there was no significant difference in the groups treated with water, ABA or iso-PhABA. The soluble acid content ranged from 0.47 to 0.54 (Figure 6c). In addition, the soluble solid content (Figure 6d and Figure 6e) and tannin content (Figure 6f and Figure 6g) in all groups treated with ABA, iso-PhABA or water showed a consistent trend, and no differences were observed among them. This shows that ABA and iso-PhABA do not affect the yield of grapes or other quality traits.

**Figure 6.**
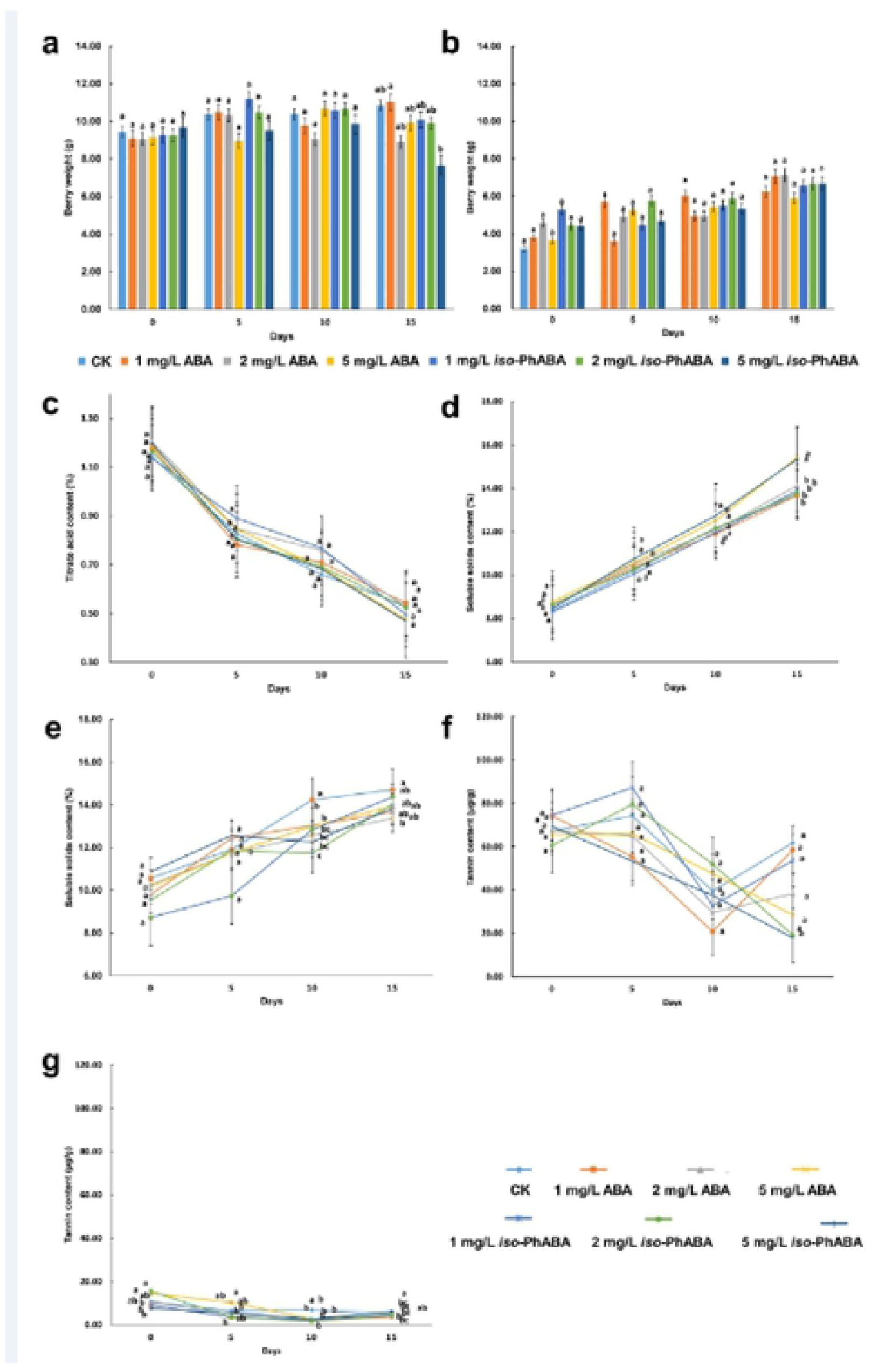
Evaluation of grape quality after treated with ABA and iso-PhABA. Weight of fruits in grape varieties Jufeng (a) and Xiahei (b); Titrate acid content of fruits in grape varieties Jufeng (c); Soluble solids content of fruits in grape varieties Jufeng (d) and Xiahei (e); Tannin content of fruits in grape varieties Jufeng (f) and Xiahei (g).

The yield and quality traits, such as the sugar and tannin contents, are important indicators in grapes, and changes in the other traits did not affect these important economic indicators. In this study, it was found that ABA or iso-PhABA treatment did not reduce the fruit weight or the contents of soluble solids and soluble acids.

## Conclusions

Grape berry color is an important factor affecting the economic value of grapes. It is mainly determined by pigments such as anthocyanins. To promote the application of ABA-like PGRs in viticulture, we selected two popular grape varieties, ‘Jufeng’ and ‘Xiahei’, and carried out a coloration experiment in Hebei and Hubei using a novel powerful ABA mimic named iso-PhABA. The results showed that in the ‘Jufeng’ and ‘Xiahei’ varieties, compared to ABA, iso-PhABA significantly increased the total content of anthocyanins at 2 mg/L and 5 mg/L. However, in these two different varieties, the trends in the anthocyanin and delphinidin contents were slightly different. The contents of anthocyanin and delphinidin in ‘Jufeng’ grapes were higher than those in the control on the 5th day after ABA or iso-PhABA treatment, and with increased time, the increase was more significant. In ‘Xiahei’ grapes, only the anthocyanin content increased significantly, and the content of delphinidin did not change significantly. This suggested that different grape varieties different responses to ABA-like PGRs. Further analysis showed that compared to ABA, iso-PhABA treatment moderately to significantly promoted the activities of some enzymes catalyzing anthocyanin. To further explore the mechanism of iso-PhABA, the affinity between iso-PhABA and ABA receptors and its inhibitory effect on the PP2C HAB1 were also investigated. The results showed that iso-PhABA showed higher affinity with some ABA receptors and displayed higher inhibition of HAB1, which might be the reason for the activity of iso-PhABA in the improvement of the color of grapes being higher than that of ABA.

Another advantage of iso-PhABA treatment is that it promotes grape coloration, similar to ABA, without any adverse effects on the quality of the grapes, such as those detected with ethephon treatment. Moreover, the soluble sugar content of ‘Jufeng’ and ‘Xiahei’ grapes treated with iso-PhABA for 10 days was similar to that of grapes treated with water for 15 days, indicating that iso-PhABA promoted the ripening of grape fruits, which enables the harvest of the treated grapes earlier than the harvest of the untreated grapes to meet market demand. It is also important to emphasize that during the phenophase of 20%-25% of the grape color change, the roots irrigated at a concentration of 2-5 mg/L of iso-PhABA racemate just once achieved or exceeded the effect of the same concentration of S-ABA. The actual effect of iso-PhABA is not only labor-saving but also more economical than spraying the ears, which can greatly reduce the application cost; thus, iso-PhABA has good application potential in promoting the coloration of grape fruit.

## Acknowledgements

This work was supported by the National Technology R&D Program in the 13th Five Year Plan of China (Grant No. 2017YFD0201303) and the national Natural Science Foundation of China (No. 21572265).

## Appendix A. Supplementary data

Supplementary material related to this article can be found, in the online version, at doi:

